# Integration of fear learning and fear expression across the dorsoventral axis of the hippocampus

**DOI:** 10.1101/2024.01.26.577384

**Authors:** Marco N. Pompili, Noé Hamou, Sidney I. Wiener

**Affiliations:** Centre Interdisciplinaire de Recherche en Biologie (CIRB) - CNRS UMR 7241 - INSERM U1050, Collège de France, Université PSL, 11 place Marcelin Berthelot, 75005 Paris, France; Institut de Neurosciences des Systèmes (INS), INSERM UMR1106, Aix-Marseille Université, 27 Boulevard Jean Moulin, 13005 Marseille, France

## Abstract

Classically, the dorsal and ventral hippocampus are thought to play distinct roles in fear conditioning, with the dorsal hippocampus primarily handling information about environmental cues and contexts, and the ventral hippocampus more involved in emotional processing. Both functions are essential for the learning and expression of conditioned fear responses, but how these processes are integrated remains largely unexplored. In this study, we simultaneously recorded single-unit activity from the dorsal and ventral hippocampus during fear conditioning to identify the neural dynamics that may underlie these processes and their integration. As fear expression emerged, shifts in neural firing patterns were observed in both regions, with a stronger shift in ventral hippocampal activity, as expected. However, contrary to the prevailing view of the ventral hippocampus as central to anxiety and fear regulation, fear expression-related neuronal responses were surprisingly more predominant in the dorsal hippocampus. In contrast, ventral hippocampal neuronal activity was more closely linked with the acquisition of conditioned fear. These features were combined in cell assemblies that emerged during fear conditioning, composed of both dorsal fear expression-responsive neurons and ventral fear learning-responsive cells. These multifactorial engrams, distributed along the hippocampal dorso-ventral axis, provide a potential substrate for integrating fear acquisition and expression, thereby coordinating associative learning.

## INTRODUCTION

The hippocampus (HPC) has long been recognized as a crucial hub for memory processes and spatial cognition^1^. While the same basic circuitry is maintained throughout the entire structure, there are anatomical, genetic and functional variations along its septo-temporal axis^2,3^. Lesions and pharmacological manipulations in the dorsal (dHPC) and ventral (vHPC) HPC in rodents (respectively, septal and temporal poles in primates) tend to have distinct impacts on spatial vs. emotional learning^4–7^. In fear conditioning paradigms, manipulations of the dHPC impair the formation, consolidation, and retrieval of contextual, but not cued fear memory^4,8,9^ while vHPC manipulations tend to have broader effects^7,10–13^. This provides support for the conventional view that the dHPC primarily processes spatial and contextual information, while the vHPC is more involved in emotional processing^2,3,14^.

This dichotomy is supported by neuronal recordings in rodents. First, the precision of spatial coding diminishes along the dorsoventral axis of the HPC^15–18^. Second, neurons in more ventral parts of the HPC exhibit a stronger association with the emotional valence of locations^19–25^ and odors^21,26^ as compared to the dHPC. Moreover, recent work further suggests that specific pathways between the vHPC, the amygdala, and the prefrontal cortex are key for anxiety and fear-related behaviors^13,27–32^.

However, this dominant view is challenged by several results highlighting the involvement of the dorsal hippocampus (dHPC) in emotional processing, beyond its traditional role in contextual or spatial processing. Indeed, under several experimental conditions, suppressing activity in the dHPC impairs trace fear conditioning^33,34^. Additionally, the modulation of fear-related dHPC engram cells impacts fear expression^35,36^, similarly to the manipulation of vHPC engram cells^37^. These results support a more nuanced view where both the vHPC and dHPC could contribute to emotional processing, and in particular to fear conditioning. Overall, while an extensive literature supports the notion that dorsal and ventral HPC play separable roles in fear conditioning, it is unknown how their relative roles are reflected in the neural activity of each subregion. Moreover, while recent literature points towards an overlap of these contributions along the hippocampal dorso-ventral axis, the mechanisms allowing such integration remain unexplored. To study this, we directly compared the activity of simultaneously recorded neurons of dHPC and vHPC in rats during Pavlovian fear conditioning. While response profiles in both dHPC and vHPC varied with fear conditioning, surprisingly, at a population level only dHPC was strongly modulated by fear expression, demonstrating that the encoding of emotionally relevant information is not restricted to the ventral hippocampus. Furthermore, more vHPC neurons developed specific responses associated with fear learning. Thus, along the dorsoventral axis of the HPC, neurophysiological representations were differentially distributed, and corresponded to complementary aspects of the emotional experience. Furthermore, during conditioning, we observed the formation of synchronous groups of neurons, i.e. cell assemblies, composed of both dHPC and vHPC members. The emergence of these assemblies provides a potential mechanism by which distinct representations of fear learning and fear expression are combined to construct an integrated representation of emotional learning across the whole hippocampus.

## RESULTS

To study the neural mechanisms underlying the distinct and shared contributions of dorsal and ventral HPC to fear learning, we simultaneously recorded in these structures in four male rats undergoing classical Pavlovian conditioning. Multiple bundles of twisted microwires were chronically implanted bilaterally in both regions, enabling the detection of single units (**Figure 1A**, **Supp. Fig. 1**, and **Supp. Fig. 2**). The behavioral protocol included two days of habituation sessions (HAB), two days of fear conditioning (FC), and one day of fear recall test (**Fig. 1B**). During these sessions, two different auditory cues were presented to the animals: CS^+^ that, during FC, co-terminated with an aversive stimulus (unconditioned stimulus, US, a mild electrical footshock), and control stimuli (CS^-^), pseudorandomly interspersed in between CS^+^ and not associated to the US. In order to study the initial acquisition of FC, we analyzed recordings from the first day of conditioning, taking the last day of HAB as a control. To measure conditioned fear, we assessed the time the animals spent freezing, using the operational definition as the periods the animals spent immobile but not asleep^39^ (see Methods). During FC, CS^+^ onsets provoked startle responses and shock delivery led to bursts of motor activity (**Supp. Fig. 3**). Therefore, most freezing events fell outside CS presentations (**Supp. Fig. 3**). Hence, to measure conditioned fear, we assessed freezing during CS^+^ intertrial intervals (ITIs; see Methods). During FC, the animals demonstrated freezing responses in the ITI after the second coupling of the CS^+^ with the US, indicating that the CS-US association was acquired then (**Fig. 1C-D**).

**Figure 1.**
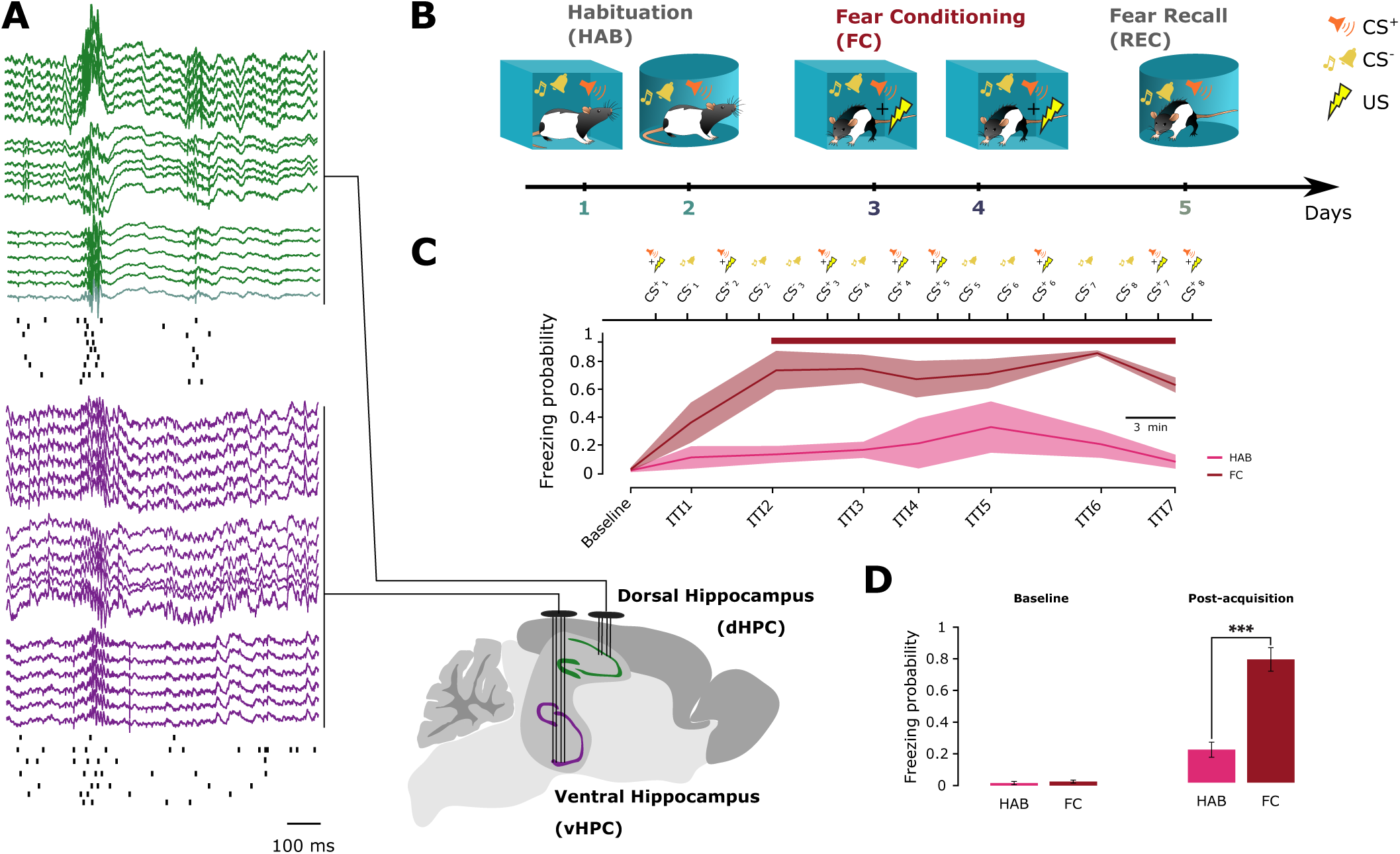
Simultaneous recordings in dorsal and ventral hippocampus during fear conditioning. (**A**) Recordings of multiple single units and local field potential in dHPC and vHPC (top) during awake immobility. Unfiltered traces appear above. Sharp wave-ripple events^38^ are accompanied by synchronous neuronal activity (below). (**B**) Behavioral protocol. On days 1 and 2, rats were habituated to the conditioning chambers and to auditory cues (CS^+^ and CS^-^) but no foot shocks (”HAB” sessions). On days 3 and 4, the rats were fear conditioned, where tone presentations were coupled with foot shocks. On day 5, correct recall of conditioned fear towards CS^+^ was tested in a chamber different from that where FC took place. (**C**) During the first fear conditioning session, the CS-US association was learned quickly. From CS2 on, the horizontal bar indicates that the freezing probability was significantly higher in the FC session than HAB session (*p <* 0.05, paired t-test). ITI: intertrial interval (the periods between successive 20-sec CS presentations). Comparison of the average amount of time spent freezing during (Left) data from from baseline HAB vs. FC sessions (prior to conditioning acquisition) and (Right) in ITI3 to 7 in HAB vs FC sessions (after conditioning acquisition).

### Shifts in hippocampal activity during fear conditioning

Previous work showed that dHPC neurons change their behavioral correlates after conditioning^40^, although no studies have reported their activity during conditioning or studied how vHPC neural activity relates to that in dHPC. To understand the respective roles of vHPC and dHPC during the actual learning process, we studied the neuronal activity in dHPC and vHPC prior to, during, and right after the acquisition of conditioning. First we examined global activity of the neuron populations, then of the individual neurons. Principal component analysis (PCA) was used to reduce the high-dimensional multi-neuronal activity of each recording session into lower-dimensional spaces. This analysis revealed that global HPC population activity shifts after the presentation of the first conditioned stimulus (CS), with a significantly greater shift during FC than during habituation (HAB) (**Fig. 2A-B**). This global shift in PCA space could reflect changes in neuronal firing rates^41^.

**Figure 2.**
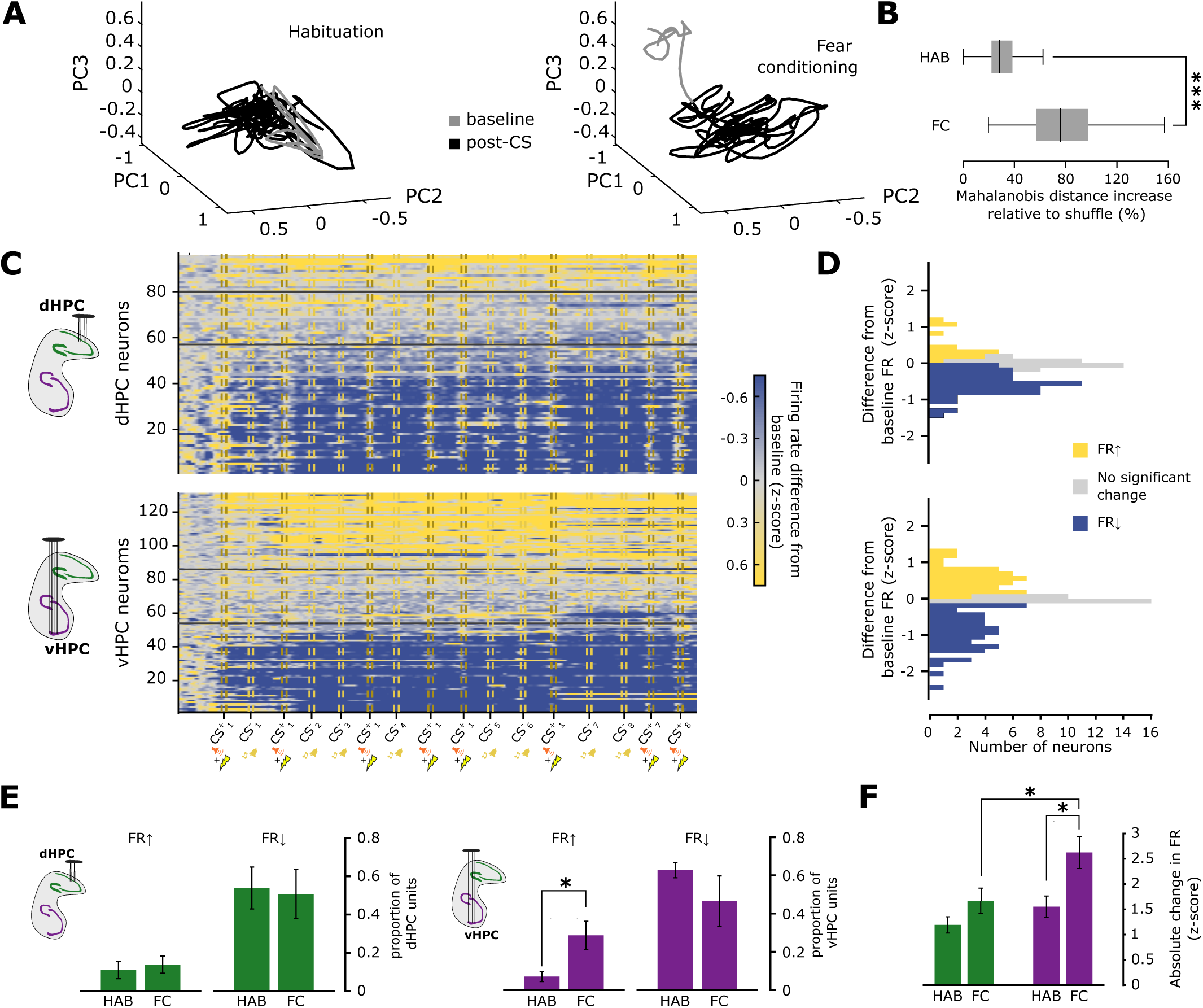
Population coding shift and neuronal firing rate changes during fear conditioning. (**A**) Evolution of PCA-projected population spiking activity over time. The first three PCA components are shown for one example animal. The gray trace corresponds to the first 3 minutes of recording before presentation of CS1, while the black trace corresponds to the period after CS^+^1. (**B**) Comparison between FC and HAB of average spatial shift (Mahalanobis distance) in PCA space between pre- and post-CS^+^1 activity relative to time-shuffled projections for all animals. (*p <* 0.001, Wilcoxon rank-sum test). (**C**) Normalized firing rates (z-scored and subtracted from baseline) during the entire FC sessions. Baseline: period before CS1. Horizontal black bars (left) separate neurons with significantly (p *<* 0.05) increasing (FR*↑*) and decreasing firing rates (FR*↓*). (**D**) Distributions of firing rate changes in A. (**E**) Comparison of relative proportions of response types. (F) Magnitudes of changes in firing rate (z-scored).

Indeed, during FC, neurons significantly changed their firing rate after the first CS, when the CS-US association had been acquired (**Fig. 2C**), with some rates increasing and others decreasing (**Fig. 2D** and **Supp. Fig. 4**). Interestingly, in the vHPC, but not dHPC, both the overall magnitudes of changes in firing rate (**Fig 2C**) and the proportion of neurons with firing rate increases (**Fig 2E**) were higher during FC than in HAB. During FC, these magnitudes were greater in vHPC than in dHPC neurons (**Fig. 2F**). These observations provide evidence for more pronounced fear-conditioning related rate coding in the vHPC than in the dHPC.

Note that the majority of HPC neurons tended to decrease their firing rate across both HAB and FC, in concert with the decrease in motor activity observed in both session types (**Supp. Fig. 5**). Indeed, as we observed previously^39^, this motor activity decrease is associated either with increases in freezing or rest during FC and HAB, respectively (**Supp. Fig. 6A**).

### Differential encoding of fear expression in dHPC and fear learning in vHPC

To gain a clearer understanding of the correlates of these population shifts, in particular whether they were behavior-related or learning-related, we then analyzed the responses of dHPC and vHPC neurons to behavioral and task-related events. Neuronal responses to freezing (fear expression) and CSs (fear conditioning) were assessed by computing the firing rate changes following the onset of these events (see Methods). Our first hypothesis was that the rate changes in vHPC after fear learning (**Fig. 2A-C**) were associated with behavioral changes related to increased freezing (**Fig. 1C** and **Fig. 2C**). Thus, we investigated neuronal responses to freezing behavior. On average, dHPC neurons reduced firing during freezing events while vHPC neurons rates did not change significantly (**Figure 3A-B**). This result failed to support our initial hypothesis, and was surprising given the ample published evidence for a role of vHPC in controlling anxiety and fear behavior^13,23,25,27,28^. A possible explanation for the dHPC freezing-associated inhibition would be related to the motor component (immobility) rather than the emotional aspect (fear) of freezing. Previous work has shown that modulation of firing rate by animals’ speed is greater in dHPC than vHPC^18^. However, here, freezing-related firing rate reductions were greater in dHPC during FC (fearful freezing) compared to freezing periods during HAB (fear-free freezing; **Figure 3C**). Thus the dHPC neural responses to freezing were not exclusively motor-related. This was further supported by the observation that while the firing rate was positively correlated with animals’ motor activity in both dHPC and vHPC (**Figure 3D**), this correlation was significantly greater in dHPC than vHPC during FC, but not during HAB (**Figure 3E**). Importantly, while freezing episodes lasted longer in FC compared to HAB, the neural response magnitudes were independent of the duration of freezing episodes (**Supp. Fig. 6B**). Overall, these results suggest that dHPC neurons respond to fearful freezing and, more generally, to movement inhibition in a fearful context.

**Figure 3.**
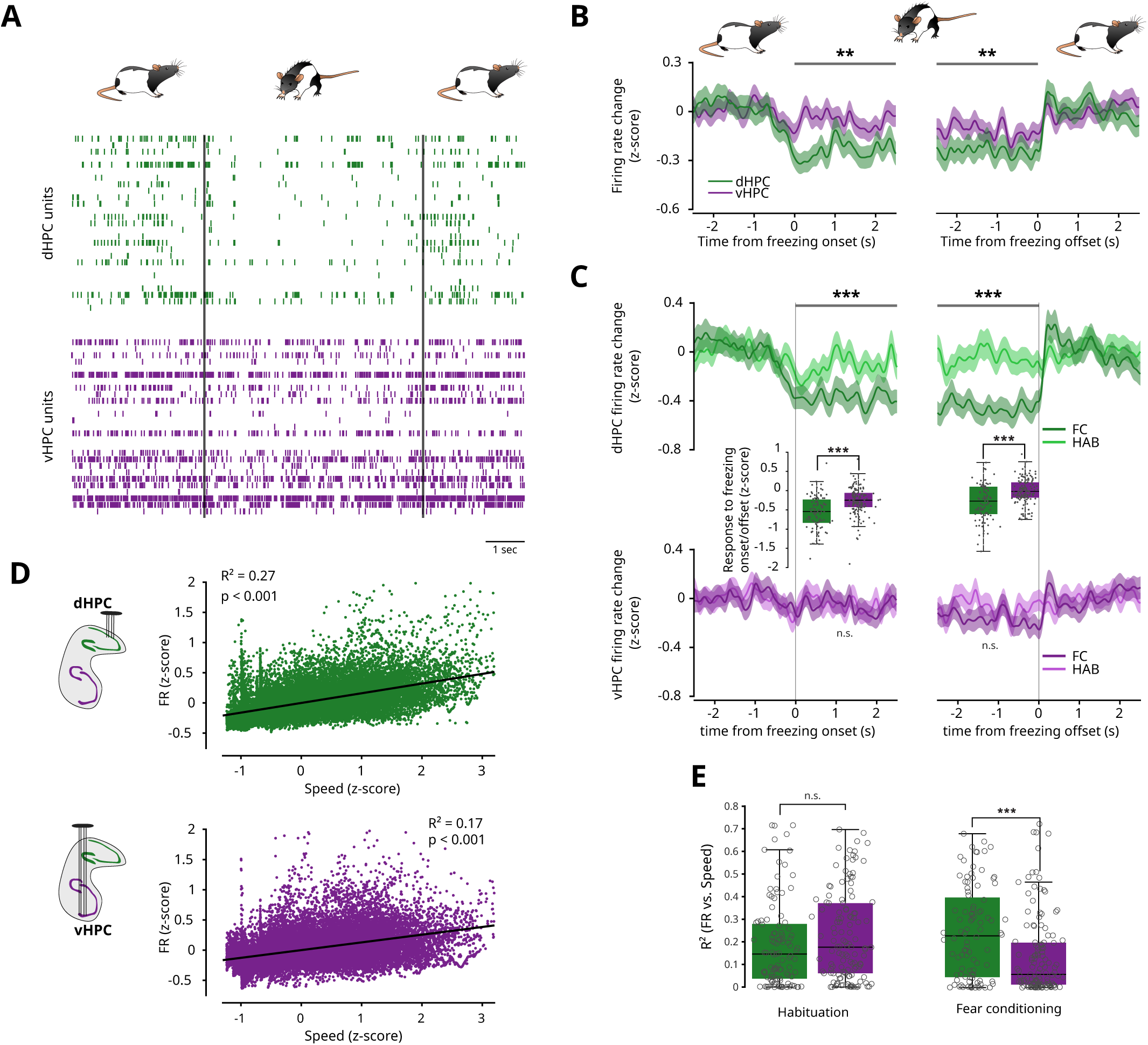
Encoding of fear expression in dHPC. (**A**) Raster plot of simultaneously recorded dHPC (blue, top) and vHPC (orange, bottom) neurons before, during and after a representative freezing episode (bracketed by vertical bars) in a FC session. (**B**) Mean (+/- SEM) firing rate changes (see Methods) in the dHPC and vHPC at freezing onset (left) and offset (right), all sessions, all animals. Bars above curves mark periods when dHPC and vHPC firing rates were significantly different (*p <* 0.01, Wilcoxon rank-sum test). Same color code as in A. (**C**) Comparison of mean firing rate changes during FC sessions (dark curves) and during HAB sessions (light curves) at freezing onset (left) and offset (right). Insets compare the vHPC and dHPC mean z-score differences after freezing episode onset and after freezing offset (see Methods) during fear conditioning (p *<* 0.001, Wilcoxon ranksum test). No significant difference in vHPC firing rate was detected between HAB and FC. (**D**) Linear regression between firing rate and animal’s speed in all data from FC and HAB sessions (1 s time bins; see Methods for their application for setting speed-correction values). (**E**) Comparison of the correlation strength (coefficient of determination) of individual neurons for linear regressions of firing rate with speed in dHPC vs. vHPC during HAB and FC (*p <* 0.001, Wilcoxon signed rank test; n.s., not significant).

In order to detect changes associated with emotional learning, we investigated neuronal responses to the CS presentations. After fear acquisition (i.e, after CS^+^2; **Fig. 1B**), neurons in both dHPC and vHPC developed excitatory responses to the CS^+^ (**Supp. Fig. 7**). To estimate the response components unrelated to movement, firing rates were corrected on the basis of the linear regression between firing rate and animals’ movement speed (**Fig. 3D-E**; **Fig. Supp. 8**, see Methods). After this correction, only vHPC neurons retained robust, sustained responses to the CS^+^ after fear acquisition (**Figure 4A-B**). Notably vHPC neurons also developed responses to CS^-^ after learning. These CS^-^ responses could be the result of fear generalization from CS^+^ to CS^-^ during FC (**Supp. Fig. 3**). Nevertheless, a closer look revealed that the responses to the individual pips composing the CS were stronger to CS^+^ than CS^-^, suggesting that the vHPC was correctly discriminating between the stimuli at a perceptual timescale. Moreover, behavioral observations confirm that animals are able to discriminate between CS^+^and CS^-^: animals displayed stronger startling during fear conditioning (FC) to CS^+^ compared to CS^-^ and increased freezing behavior during fear recall to CS^+^ compared to CS^-^ (**Supp. Fig. 3**). The emergence of these vHPC CS^+^ neural responses occurred upon the acquisition of the CS-shock association (**Supp. Fig. 9**) and the observed differences in neuronal response magnitude after learning were not attributable to neurons from a single animal, since the proportion of CS^+^-responsive neurons in the vHPC was consistently higher than both the proportion of CS^-^-responsive neurons in the vHPC and the proportion of CS^+^-responsive neurons in the dHPC (**Figure 4C**). Collectively, these results support that the vHPC, but not the dHPC, plays a role in representing the learned relationship between a specific conditioned stimuli and the associated outcome.

**Figure 4.**
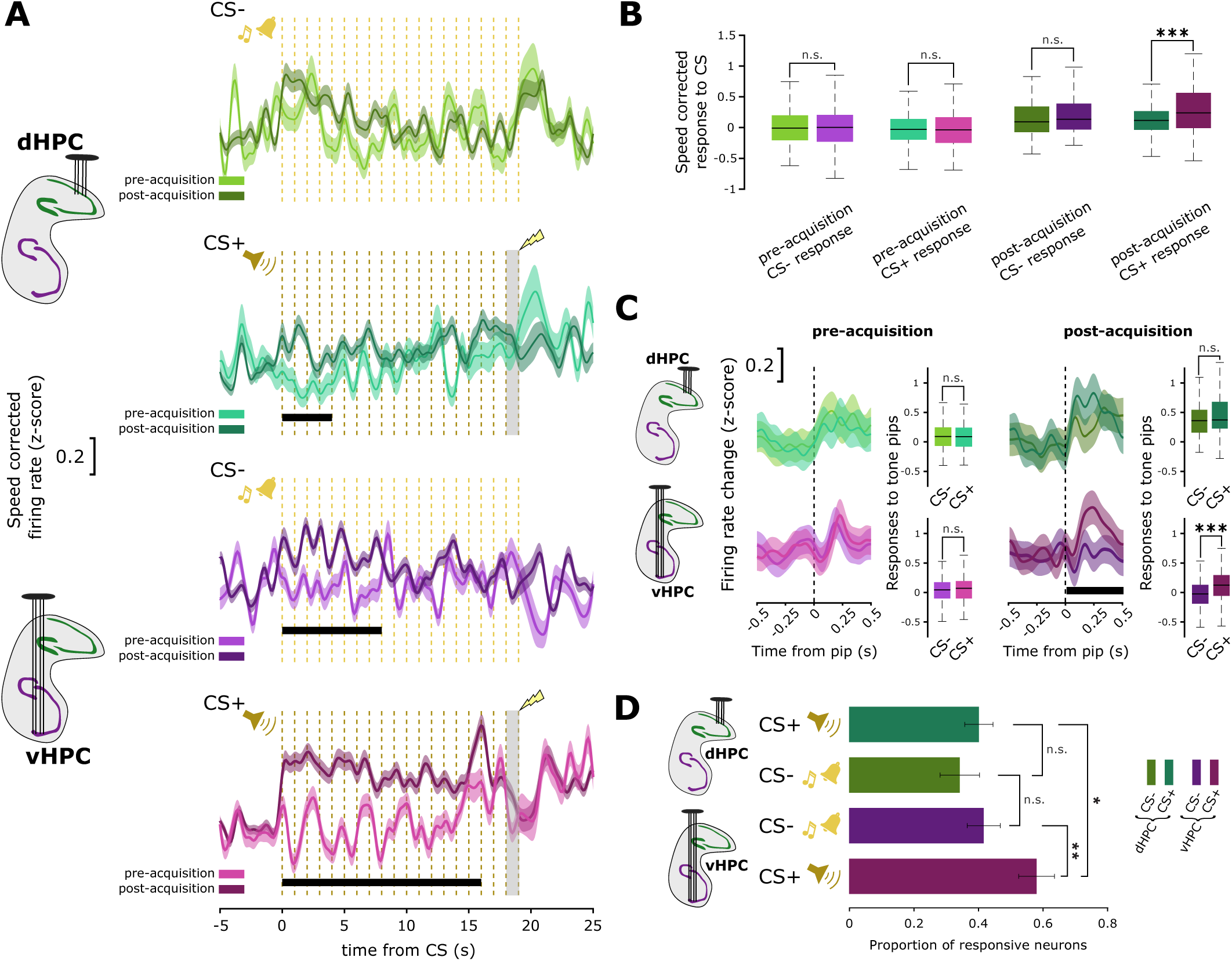
Encoding of fear learning in the vHPC. (**A**) Comparison of neurons’ speed-corrected responses to CS^+^ and CS^-^ before fear acquisition (CS^+^1 and CS^+^2, light curves) and after (CS^+^3 to CS^+^8, dark curves) during the first FC session (*p <* 0.001, Wilcoxon signed rank test). Dotted vertical bars represent the tone pips, and the gray shaded bar is the US presentation. (**B**) Comparison of the average responses to CS^+^ and CS^-^ presentations (first 10 seconds after the CS onset) of dHPC vs. vHPC, pre-acquisition and post-acquisition during the FC session (*p <* 0.001, Wilcoxon rank-sum test). (**C**) Comparison of neurons’ speed-corrected responses to the individual pips of CS^+^ vs. CS^-^, before and after fear acquisition during the FC session (*p <* 0.001, Wilcoxon signed rank test, left). Dotted vertical bars represent the pips. Right, comparison of the average responses to the pips of CS^+^ vs. CS^-^ (*p <* 0.001, Wilcoxon rank-sum test). Comparison of the proportions of neurons displaying a significant response to CS in dHPC and vHPC.

### Mutual contribution of dHPC and vHPC to mixed cell assemblies emerging during FC

Finally, we tested whether the respective response profiles in the dHPC and vHPC during FC are coordinated, potentially integrating representations of fear expression and fear learning. For this, we examined the synchronization of neurons across the HPC dorso-ventral axis in each recording session by extracting cell assemblies from PCA components (**Fig. 2A-B**) using ICA^42–44^. We found assemblies composed of only dHPC neurons (dHPC-only), composed only by vHPC cells (vHPC-only) and *mixed assemblies*^43^, composed of neurons from both the vHPC and the dHPC (**Figure 5A** and **Supp. Fig. 9**). Activity rates of vHPC-only and mixed assemblies changed significantly after the first tone-shock association, while dHPC-only assemblies did not show a significant response (**Figure 5B-C**). Moreover, while there was a low incidence of mixed assemblies during HAB, their proportion was significantly higher during FC. In contrast, the proportion of vHPC-only assemblies decreased (**Figure 5D**). This would be consistent with vHPC assemblies recruiting dHPC neurons during FC to form mixed assemblies, possibly integrating signals from both areas. Consistently, while dHPC-only assembly neurons primarily responded to freezing, and vHPC-only assembly neurons to conditioning, mixed assemblies were composed of both dHPC neurons responding to freezing and vHPC neurons responding to the acquisition of conditioning (**Fig. 6A-D**). This suggests that fear learning and expression representations can be integrated through dorso-ventral intrahippocampal synchronization (**Fig. 6E**).

**Figure 5.**
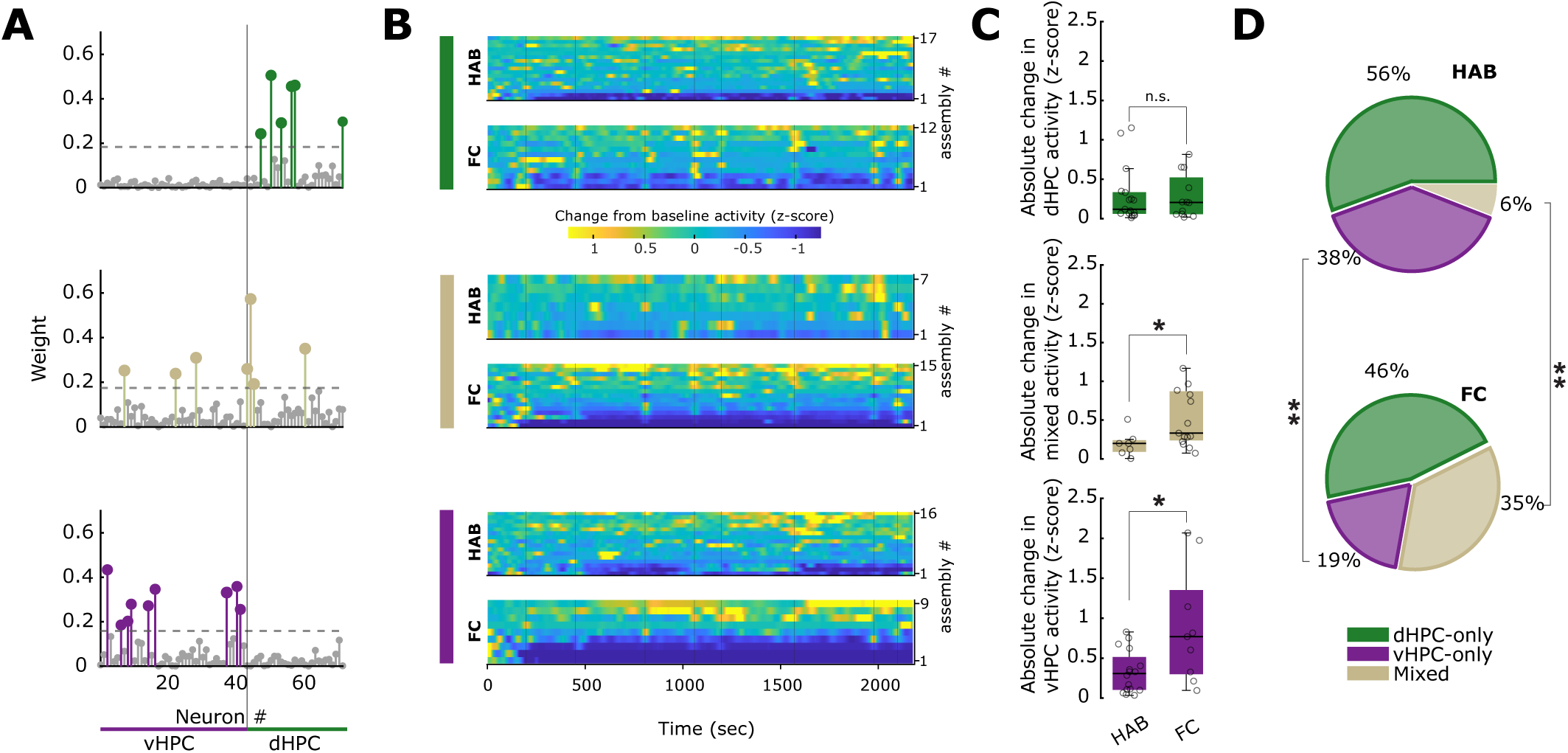
Recruitment of mixed dorsal-ventral assemblies during fear conditioning. (**A**) Example dHPC-only (top), mixed (middle) and vHPC-only (bottom) assemblies. Each circle corresponds to one neuron. Large colored circles correspond to assembly members and small gray circles to non-members. The lengths of the bars depict each neuron’s contribution (ICA weight) to the assembly. Horizontal dashed lines represent the threshold for a cell to be considered a member of the assembly (see Methods). (**B**) Activity of the dHPC-only (top), mixed (middle), and vHPC-only (bottom) assemblies during FC and HAB sessions. Vertical bars correspond to CSs occurences. (**C**) Absolute activity changes of assemblies after CS^+^1 relative to baseline. (**D**) Average proportions of dHPC-only, vHPC-only and mixed assemblies. Brackets indicates significant differences between proportions (*p <* 0.01, paired t-test). All except mixed HAB are significantly greater than zero (*p <* 0.01).

**Figure 6.**
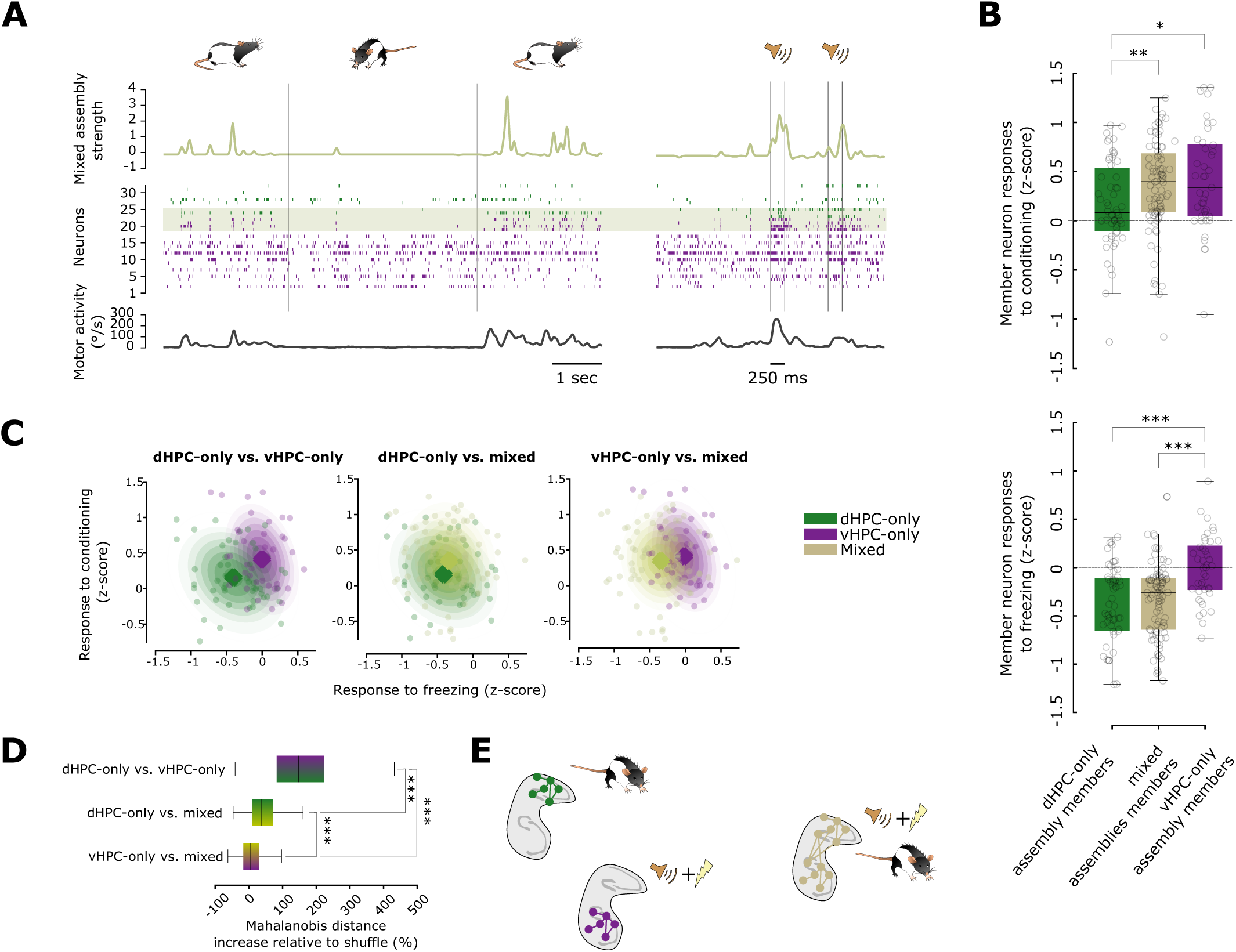
Mixed dorsal-ventral assemblies are composed of dHPC fear expression responsive neurons and vHPC fear learning responsive neurons. (**A**) Representative example showing assembly activation strength (top), raster plots of single unit activity (middle), and animal’s speed (bottom) during (left) a freezing episode (bracketed by vertical bars), and (right) during CS^+^ presentations in FC1 after acquisition (pips onsets and offsets are bracketed by pairs of vertical bars). vHPC unit spikes are in green and dHPC in purple. The rows with the gray shading correspond to the assembly members. (**B**) Comparison of the average responses to freezing and conditioning for members of dHPC-only, mixed, and vHPC-only assemblies. Same data points of Figures 2C and 3A but grouped according to assembly composition. (**C**) Comparison of the responses to freezing and conditioning of the member neurons grouped by assembly types’. Dots: responses of each assembly member. Large dots correspond to centers of mass. Shaded areas, contour lines of Gaussian model density estimates of the clusters’ distributions. (**D**) Comparison of the distances increases between the clusters in the plots in G relative to clusters with shuffled neuron identities. (**E**) Schematic representation of dHPC-only assembly neurons responding to freezing, vHPC-only assembly neurons to conditioning, and mixed assembly neurons to fearful freezing and fear conditioning.

## DISCUSSION

Multiple decades of research have demonstrated, with lesion, pharmacological, and optogenetic approaches, that dHPC and vHPC play different roles during emotional learning, mainly processing contextual and emotional information, respectively. More recent findings challenge the strict dichotomy of earlier models, suggesting that, despite their specificities, dHPC and vHPC work in concert to sustain adaptive behavior. Nevertheless, simultaneous recordings of single units in these regions had not been reported before, and it was unknown how this integration is implemented or even whether neural populations in these hippocampal regions express distinct activity profiles.

Here we show that the activity profiles of populations of neurons of both dHPC and vHPC underwent significant shifts during fear conditioning, with vHPC exhibiting the more reorganization. Concurrently with the acquisition of the CS-US association, these shifts are associated with dHPC neurons being predominantly inhibited during fearful freezing, while vHPC neurons developed stronger excitatory responses to fear-associated CS presentations. Neurons with these distinct responses are recruited during fear conditioning by mixed dorsal-ventral assemblies of synchronous cells. This provides a potential mechanism for integrating responses of individual cells within multifactorial hippocampus-wide engrams representing both emotional learning and its associated behavioral response.

Functional differences between dHPC and vHPC in fear learning have been posited for more than three decades. However, to date, it was still unknown if and how neurons across the HPC dorsoventral axis may differentially encode fearful stimuli and behavior. Here, to our knowledge, we show for the first time, encoding differences between dHPC and vHPC during fear learning. Distributed cell assemblies spanning multiple structures were recently described across the cortex and striatum^43^. Here, exploiting simultaneous recordings of dorsal and ventral hippocampus, we were able to show that, similarly, neurons synchronize at different points along the hippocampal dorsoventral axis upon learning supporting a conjunctive representation of fear expression and fear learning. It must be noted that some vHPC-only and dHPC-only assemblies could be part of larger mixed assemblies with other members that were not detected. Following our work, a more recent report showed how mixed dorso-ventral mixed assemblies reactivate during non-REM sleep, possibly sustaining the selective reactivation of aversive experiences during sleep^45^.

Here, consistent with the standard procedure fear conditioning research in rodents, we employed freezing behavior as a measure of conditioned fear. It must be noted that automatic detection of freezing can only be measured by assessing when the animal is immobile, but that certain immobility states may not correspond to actual freezing^39^. In particular, in our protocol, many freezing episodes in HAB likely corresponded to rest, as we show elsewhere^39^. This allowed us to show that dHPC neural activity is more modulated by actual fearful freezing during FC than to the freezing detected during HAB that is largely contaminated by fear-free resting episodes.

Crucially, while past studies only compared neural activity before vs. after conditioning^24,40^, here we were able to record neural activity during FC itself. This enabled the direct study of how neural activity changes with learning, and to show that HPC representations rapidly adapt their coding properties after a single presentation of the CS-US association. This suggests a potential mechanism underpinning the acquisition of conditional fear memory after a single CS-US pairing^46^.

Previous work in trace fear conditioning showed that dHPC neurons develop a response to the conditioned stimulus (CS) but that this activity is not persistent enough to bridge the ‘trace’ delay period^47^. Similarly, in our study, we found that dHPC responses are not sustained, whereas vHPC responses are. Future work could examine whether the vHPC generates a representation that is sufficiently persistent to encode the trace period.

Anatomical connectivity gradients could explain the different responses observed in the vHPC and dHPC. The vHPC has stronger bilateral connections with the amygdala^13,27,28,32^ than the dHPC, and the amygdala is the first structure to encode the CS-US association during FC^48^. Consistently, we observed the acquisition of excitatory responses to the CS in the vHPC, and an increase in the proportion of firing rate increasing cells in the vHPC during fear conditioning. Anatomical gradients in the HPC might therefore underlie ‘representational gradients’, along which different features of experience are processed. As suggested in a recent study, dHPC and vHPC circuits could then broadcast information about these different features to distinct downstream brain structures to support memory^49^.

Another possibility is that there is a balance of dHPC/vHPC activity that is altered both during freezing and CS presentations, via a decrease of dHPC activity during freezing and via an increase of vHPC firing during CSs. Indeed, the dHPC responses to freezing are largely decreases in firing rate, suggesting that the dHPC may go through a general inhibition during fearful freezing, perhaps as a result of neuro-modulation and/or a reduction of excitatory input. Accordingly, a recent report shows how activity decreases of neural ensembles may serve neural encoding^50^.

The acquisition of a fearful freezing-specific response in dHPC neurons challenges the notion that the dHPC is restricted to spatial and contextual processing, but is consistent with a recent report showing that optogenetic stimulation of dHPC, but not vHPC during FC increases freezing^51^. These results extend a new perspective: the dHPC could encode a more comprehensive representation of the animal’s state relative to its environment. This expanded representation would not only include spatial but also emotional states, such as fearful freezing states. Conversely, the vHPC may predominantly represent external environmental contingencies, such as the CS-US association. To be able to respond to an imminent threat, the brain needs to rapidly bind these distinct representations together. Here, these complementary representations emerge rapidly during fear conditioning, after only one or two CS-US presentations, and are combined in mixed assemblies. Our study thus provides insights into the respective representations emerging in the dHPC and vHPC during fear conditioning and underscores their collective role in orchestrating an adaptive response. Future work may test whether the activation of these representations play a causal role in controlling behavior and learning.

## ACKNOWLEDGMENTS

We thank Gabriel Makdah and Eulalie Leroux for help with data acquisition, Ralitsa Todorova for fruitful discussions, and Céline Drieu and Kishore Kuchibhotla for useful comments on earlier version of this manuscript. This work was funded by a Labex MemoLife grant to SIW and MNP.

## AUTHOR CONTRIBUTIONS

Research project design: MNP, NH. Experiments: MNP. Data analysis design: MNP, NH, SIW. Data analysis: NH. Writing and results visualisation: MNP, NH, SIW

## METHODS

### Animals

Four male Long-Evans rats (350–400 g at the time of surgery) were housed individually in monitored conditions (21°C and 45% humidity) and maintained on a 12h light – 12h dark cycle. In order to avoid obesity, food was restricted to 13–16 g of rat chow per day, while water was available ad libitum. To habituate the rats to human manipulation, they were handled each workday. All experiments conformed to the approved protocols and regulations of the local ethics committee (Comité d’éthique en matière d’expérimentation animale Paris Centre et Sud n°59), the French Ministries of Agriculture, and Research.

### Surgery

The rats were deeply anesthetized with ketamine-xylazine (Imalgene 180 mg/kg and Rompun 10 mg/kg) and anesthesia was maintained with isoflurane (0.1-1.5% in oxygen). Analgesia was provided by subcutaneous injection of buprenorphine (Buprecaire, 0.025 mg/kg) and meloxicam (Metacam, 3 mg/kg). The animals were implanted with a custom-built prosthesis with 42 independently movable hexatrodes (bundles of 6 twisted tungsten wires, 12 *µ*m in diameter, gold-plated to ∼200 kΩ). The electrode tips were implanted 0.5 mm above the CA1 pyramidal layer of dHPC and vHPC −5 to −4.5 mm posterior and +/- 3.8 to 5 mm lateral from bregma. Miniature stainless steel screws were implanted above the cerebellum to serve as electrical reference and ground. During recovery from surgery (minimum 7 days), the rats received antibiotic (Marbofloxacine, 2 mg/kg) and analgesic (Meloxicam, 3 mg/kg) treatments via subcutaneous injections and were provided with food and water ad libitum. The recording electrodes were then progressively lowered until sharp wave ripple and single unit activity was visible and adjusted to optimize yield and stability of single-units.

### Behavioral apparatus

Two environmental contexts (BOX and CYL) were used for the fear protocol. The BOX context was a cubicle conditioning chamber (40×40×40 cm) with gray PVC walls lined with ribbed black rubber sheets and a floor composed of nineteen stainless steel rods (0.48 cm diameter with 1.6 cm intervals) connected to a scrambled shock generator (ENV-414S, Med Associates, USA). It was scented daily with mint cleaning solution (Simple Green, Sunshine Makers). The CYL context was a stadium-shaped enclosure (30 cm straight sides and 15 cm radius) with a black wooden floor, and walls lined with light brown pieces of rope rug. It was scented daily with a vanilla extract solution. A custom-made electronic system presented the animals with the two auditory CSs types (80 dB, 19.25 s long, each composed of 20 pips of either white noise, CS+, or 8 kHz pure tones, CS-, presented at 1 Hz and 250 ms long). During each session, after a baseline period of 3 min, the animals were presented to 16 CS (8 CS^+^ and 8 CS^-^), separated by random-duration intervals of silence ranging between 120 and 240 s.

### Behavioral protocol

Each experimental day, rats were placed in both contexts for 35 min to ensure that any behavioral differences observed between the two environments were not related to their familiarity. Before and after recording sessions, animals were allowed to rest in a cloth-lined plastic flowerpot (30 cm upper diameter, 20 cm lower diameter, 25 cm high). On days 1 and 2, rats underwent habituation sessions in both contexts, with and without CS^+^ and CS^-^ presentations. On day 1, the CSs were presented in one of the two contexts, and on day 2 in the other. On days 3 and 4, the rats were fear conditioned in the BOX where CS^+^ presentations were coupled with foot shocks (1 s, 0.6 mA, co-terminating with the last CS^+^ pip).

### Data acquisition and processing

Brain activity was recorded with a 256-channel digital data acquisition system (KJE-1001, Amplipex, Szeged, Hungary). The signals were acquired with four 64-channel headstages (Amplipex HS2) and sampled wideband at 20 kHz. An inertial measurement unit sampled the 3D angular velocity and linear acceleration of the head at 300 Hz. To determine the instantaneous position of the animal, a red LED mounted on the headstage was imaged by overhead webcams at 30 Hz. Animal behavior was also recorded at 50 Hz by two lateral video cameras (acA25000, Basler) in each environment. Off-line spike sorting was performed using KlustaKwik (K.D. Harris, http://klustakwik.sourceforge.net). The resulting clusters were visually inspected using Klusters^52^ to reject noise and to merge units erroneously considered as different units. Neurophysiological and behavioral data were explored using NeuroScope^52^. LFPs were derived from wideband signals by downsampling all channels at 1250 Hz.

### Scoring of behavioral and brain states

Automatic detection of immobility was performed by thresholding the angular speed calculated from gyroscopic data as described in^53^, where we showed that measurements of angular velocity are more suitable than those of linear acceleration for discriminating motion from immobility. LFP data was visualized using Neuroscope^52^ and sleep and freezing were detected as previously described^39^. CS^+^ ITIs were defined as the time intervals between the presentations of two CS^+^ and deducted from the presentation time of CS^-^.

### Histological identification of recording sites

At the end of the experiments, recording sites were marked with small electrolytic lesions (∼20 μA for 20 s, one lesion per bundle). After a delay of at least three days to permit glial scarring, rats were deeply anesthetized with a lethal dose of pentobarbital, and intracardially perfused with saline (0.9%) followed by paraformaldehyde (4%). Coronal slices (35 μm) were stained with cresyl-violet and imaged with conventional transmission light microscopy. Recording sites were reconstructed by comparing the images with the stereotaxic atlas of^54^. Data recorded in electrodes bundles whose location could not not be identified were discarded from further analyses.

### Statistical analysis

All statistical analyses were performed in Matlab (MathWorks, Natick, MA), using the Freely Moving Animal Toolbox (http://fmatoolbox.sourceforge.net) and custom written programs. For descriptive statistics, behavioral data was represented with boxplots where the central bar indicates the median, the bottom and top edges indicate the 25th and 75th percentiles; whiskers extend to the most extreme data points, excluding outliers. Datapoints were considered as outliers if they were greater than *q*3 + 2.7*σ*(*q*3–*q*1) or less than *q*1–2.7*σ*(*q*3–*q*1); where *σ* is the standard deviation, and q1 and q3 are, respectively, the 25th and 75th percentiles of the sample data.

### Analysis of population trajectories over time

Principal Component Analysis (PCA) of figure 1D was computed on the spike trains of all hippocampal neurons for each rat. The bins corresponding to shocks and the following 10 seconds were removed in order not to bias population activity with the strong shock responses. For each rat, we measured the distance of projections occurring before and after the first CS^+^ presentation and the distance of temporally shuffled projections and computed the % increase in distance between both distance measures. The distance metric used here was the Mahalanobis distance. Given two points defined by the coordinates **x** and **y** and the covariance matrix, the Mahalanobis distance between these points is defined as:

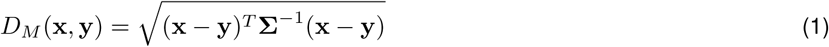

### Assessment of freezing and conditioning responses

Firing rate responses to freezing events and CS presentations were assessed by analyzing their peri-event firing rate changes. This was computed by zscoring the raw firing rate and subtracting the average baseline firing rate. Baseline was 2.5 s before freezing onsets for freezing responses and 10 s before CS presentations for CS responses. CS periods were not excluded from the analyses of freezing events and vice-versa. In order to assess how CS responses changed with learning, we computed the learning induced change of the responses as the average firing rate change in the 10 seconds following CS presentation (as shown in figure 2F) normalized by subtracting the mean response to the first CS (**Supp. Fig. 9**).

### Identification of freezing-responsive neurons

To identify neurons that are freezing-responsive, we compared the average z-scored firing rate during the first 10 seconds after the sound onset before learning (CS 1 and 2) vs after learning (CS 3 to 8). If the p-value (Wilcoxon signed-rank test) was below 0.05 and the change in z-score was positive, the neuron was identified as freezing-responsive.

### Analysis firing rate vs. speed relationship

In order to study the relationship between neurons firing rate and animals’ motor activity we performed linear regressions. For each neuron we performed a linear regression between speed and the firing rate (z-scored and downsampled in 1 s bins) and computed the coefficient of determination. This was performed individually for each neuron with all time bins of all session types. The distributions of the coefficient of determination in dHPC vs. vHPC were then compared.

### Correction for speed-related firing rate changes

To correct for speed-related changes in firing rate, for each neuron, we performed a linear regression between firing rate and the angular velocity of the head of the animal on the CS-presentation window. We then removed the linear component of the regression from the firing rate of the neuron according to the animal’s speed at that time. The angular velocity of the head was used because it is a more precise measurement of body movement then linear velocity^53^. While the gyroscopes of the headstages measuring the head angular speed provide improved precision for low speed states compared to measurements of body linear velocity, it is important to note that when the animal is walking/running both angular and linear velocity are characterized by high speed.

### Identification of cell assemblies

A standard unsupervised method based on principal and independent component analyses (PCA and ICA) detected the co-activation of simultaneously recorded neurons^55^ as we described previously^43,44^. Briefly, spike trains were first binned into 15-ms bins and z-scored to generate a z-scored spike count matrix Z, where *Z_i,j_* represents the activity of neuron i during time bin j. The same removal of shock-related bins was performed as in the PCA. Principal components (PCs) were computed by eigendecomposition of the correlation matrix of Z. Principal components associated with eigenvalues exceeding the upper bound of the Marchenko-Pastur distribution were considered significant. We then carried out ICA (using the fastICA algorithm of H. Gavert, J. Hurri, J. Sarela, and A. Hyvarinen, http://research.ics.aalto.fi/ica/fastica) on the projection of Z onto the subspace spanned by significant PCs. Independent component (IC) weights were scaled to unit length, and by convention the arbitrary signs of the weights were set so that the highest absolute weight was positive. Members of cell assemblies were identified using Otsu’s method to divide the absolute weights into two groups maximizing inter-class variance, and neurons in the group with greater absolute weights were classified as members. Goodness of separation was quantified using Otsu’s effectiveness metric, namely the ratio of the inter-class variance to the total variance. This procedure yielded a set of vectors *C_i_* representing the detected cell assemblies. We verified that members of detected assemblies were more synchronous than non-member neurons (**Supp. Fig. 10**).

### Assembly activations

We computed an instantaneous assembly activation strength as described previously^56,57^:

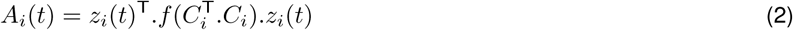

where *C_i_*is a vector of size *ν* (*ν* is the number of neurons) containing the weights of the members of the *i^th^* assembly, and *z_i_*(*t*) is a vector of size *q* (*q* is the number of time bins) of the activity of the assembly members at time *t*, and 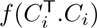 is a transformation of the outer product where the diagonal is set to 0, so that spiking in a single neuron does not contribute a high activation strength.

## SUPPLEMENTARY FIGURES

**Supplementary Figure 1.**
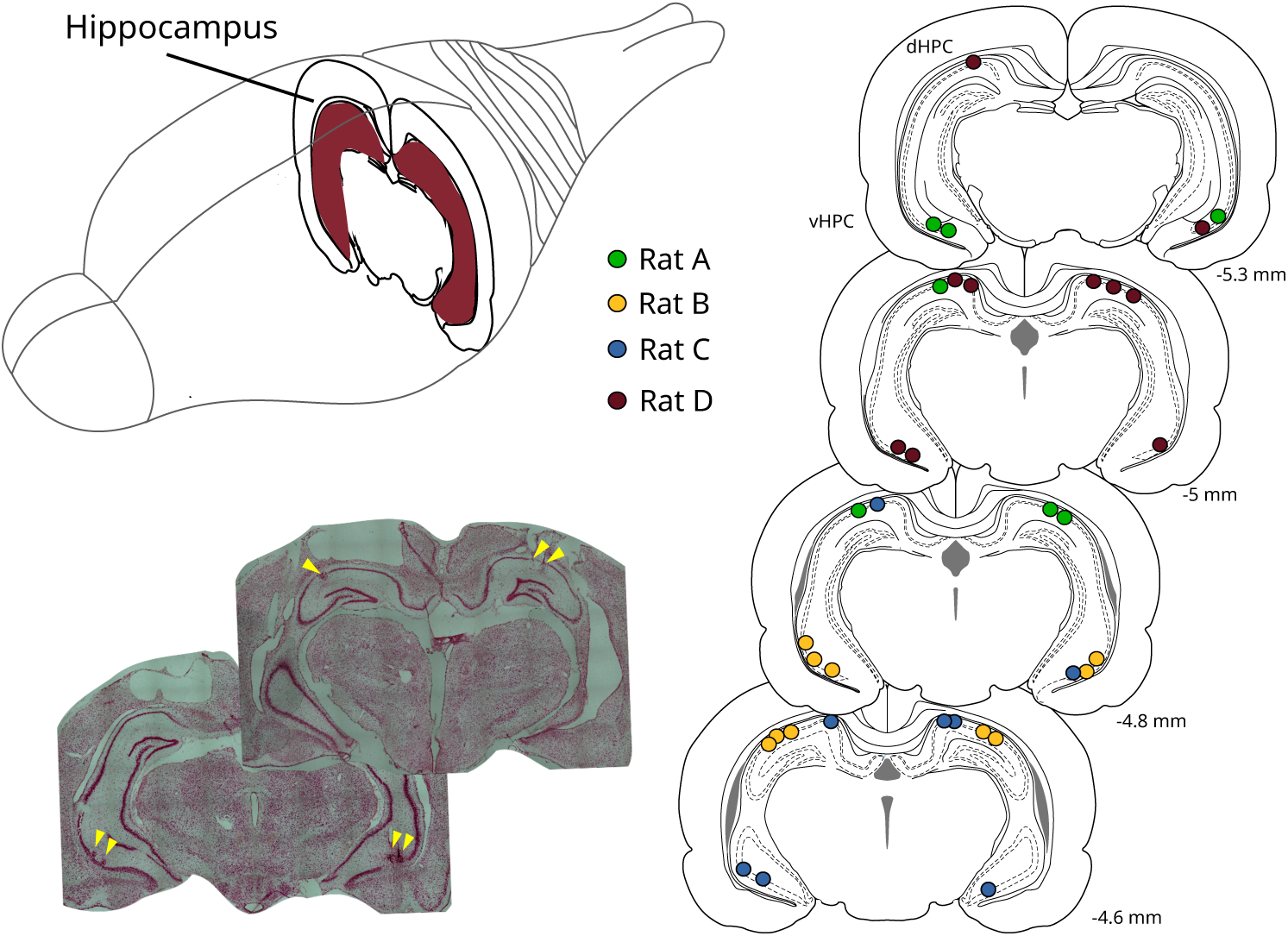
Anatomical reconstruction of the recording sites. Sites of electrolytic lesions in the four rats are indicated by the colored circles on the diagrams (right) while the arrows on the photomicrographs (bottom left) depict representative recording locations in rat B.

**Supplementary Figure 2.**
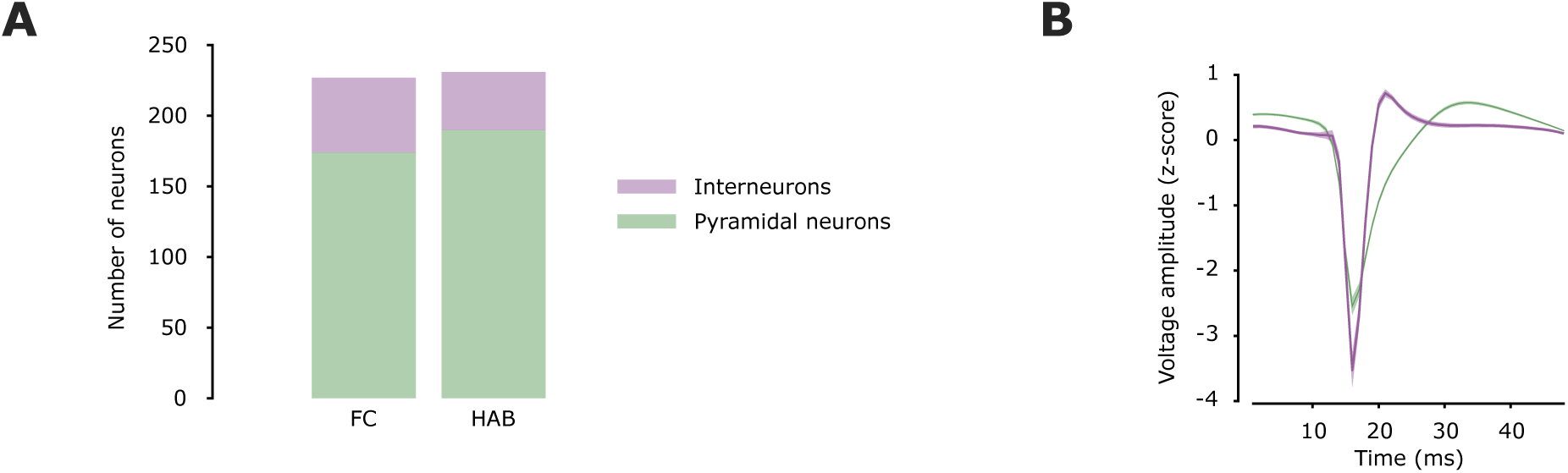
Discrimination and quantification of putative interneurons and principal neurons. (**A**) Numbers of interneurons and pyramidal neurons recorded in the FC and HAB sessions (**B**) Average waveforms for all cells in all rats (same color code as in A, shading represents SEM).

**Supplementary Figure 3.**
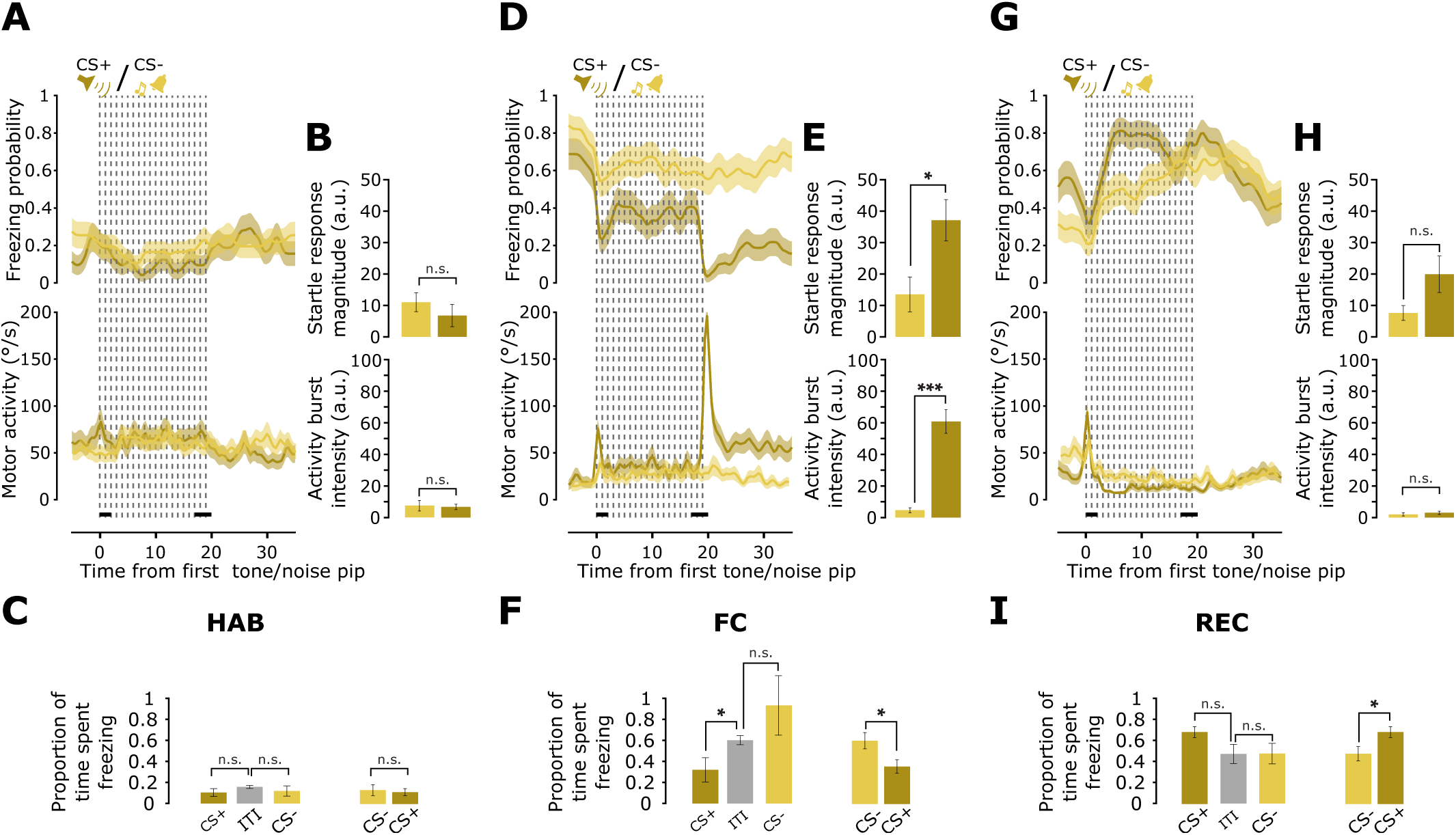
Behavior in relation to CS presentations. (**A**) Average (+/- SEM) freezing probability (top) and angular speed (bottom) during CS^+^ (dark yellow) and CS^-^ (light yellow) presentations during HAB. Dashed vertical lines represent the individual pips composing the CS. (**B**) Comparison between CS^+^ and CS^-^ of the magnitude of the startle triggered by the onset of the first pip (top) and the intensity of the motor activity burst that is triggered by the delivery of the US (bottom). These are computed with the average response during respectively 0-2 (first horizontal black line) and 17-20 seconds from the CS onset (second horizontal black line). (**C**) Comparison of the freezing rate during CS presentations and the inter-trial intervals (ITIs) during HAB. (**D**) Same as A but for FC. Note how CS onset induces a startle response and another occurs at the end of the CS (co-terminating with the US delivery). (**E**) Same as B but for FC. (**F**) Same as C but for FC. (**G,H,I**) Same as A,B and C but for REC.

**Supplementary Figure 4.**
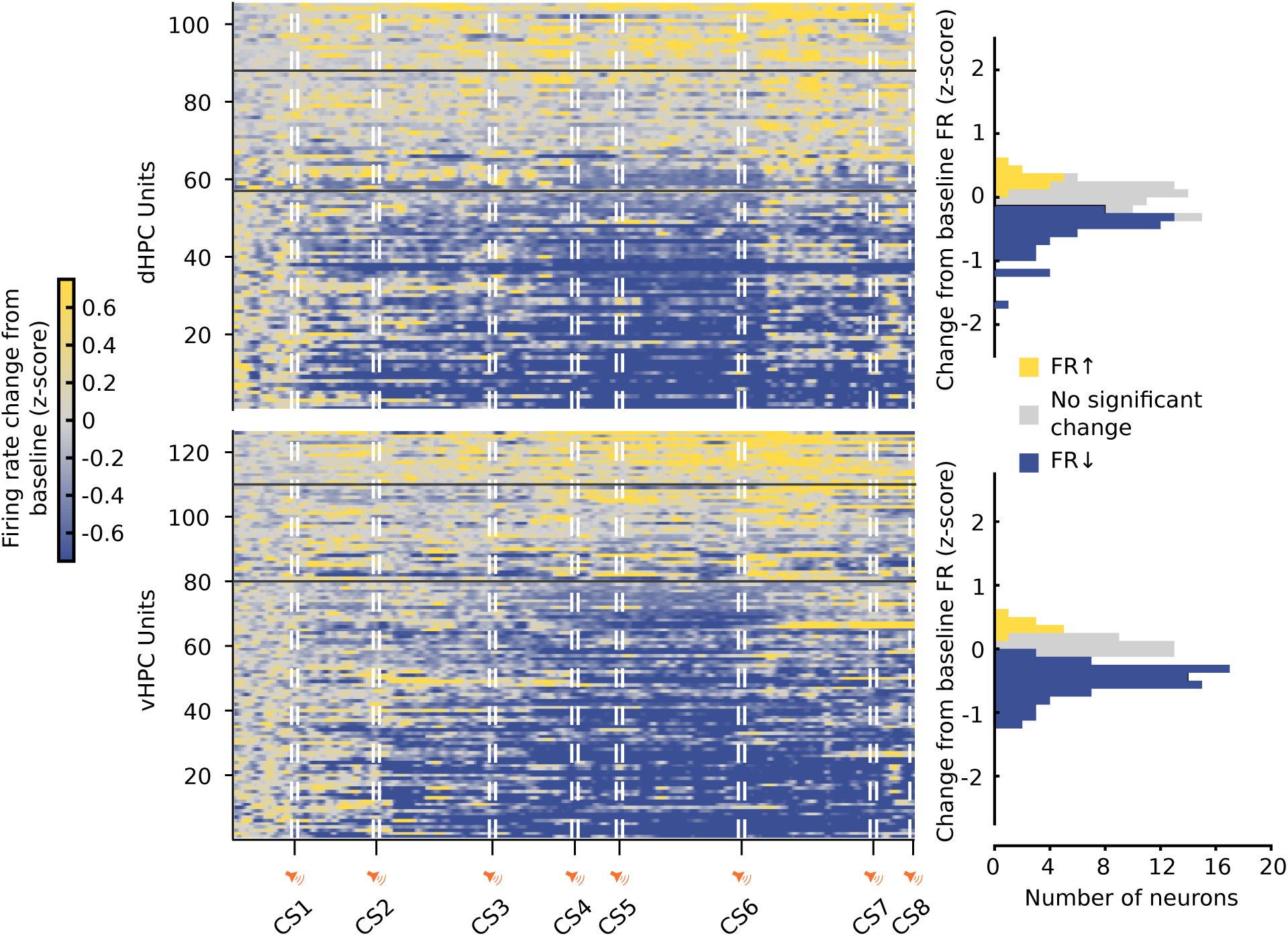
Neuronal firing rate changes during habituation sessions. Color plot of normalized firing rate (z-scored and subtracted relative to baseline) of all neurons during the second HAB session. Baseline: period before the first CS. Horizontal black bars in color rasters separate neurons with significantly (p *<* 0.05) increasing (FR*↑*), no significant change, and decreasing firing rate (FR*↓*). Histograms (right) show the distribution of magnitudes of the firing rate changes.

**Supplementary Figure 5.**
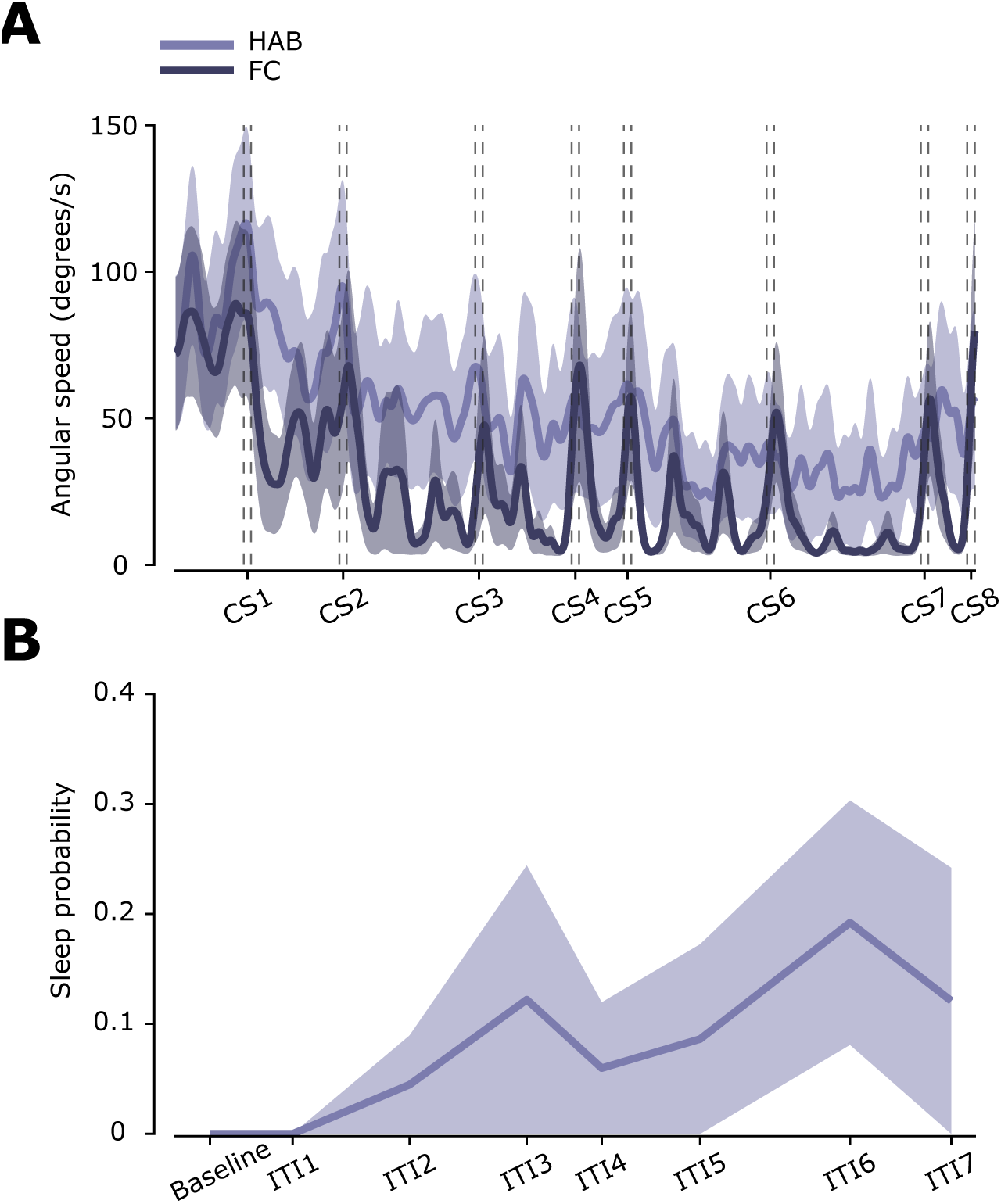
Motor activity changes over the course of the session. (**A**) Average speed during HAB (light purple) and FC sessions (dark purple). Shading corresponds to SEM. Peaks at CS presentation correspond to startle responses to the CS and to motor responses to the shock. (**B**) Average sleep probability during the HAB. No sleep periods were detected during FC.

**Supplementary Figure 6.**
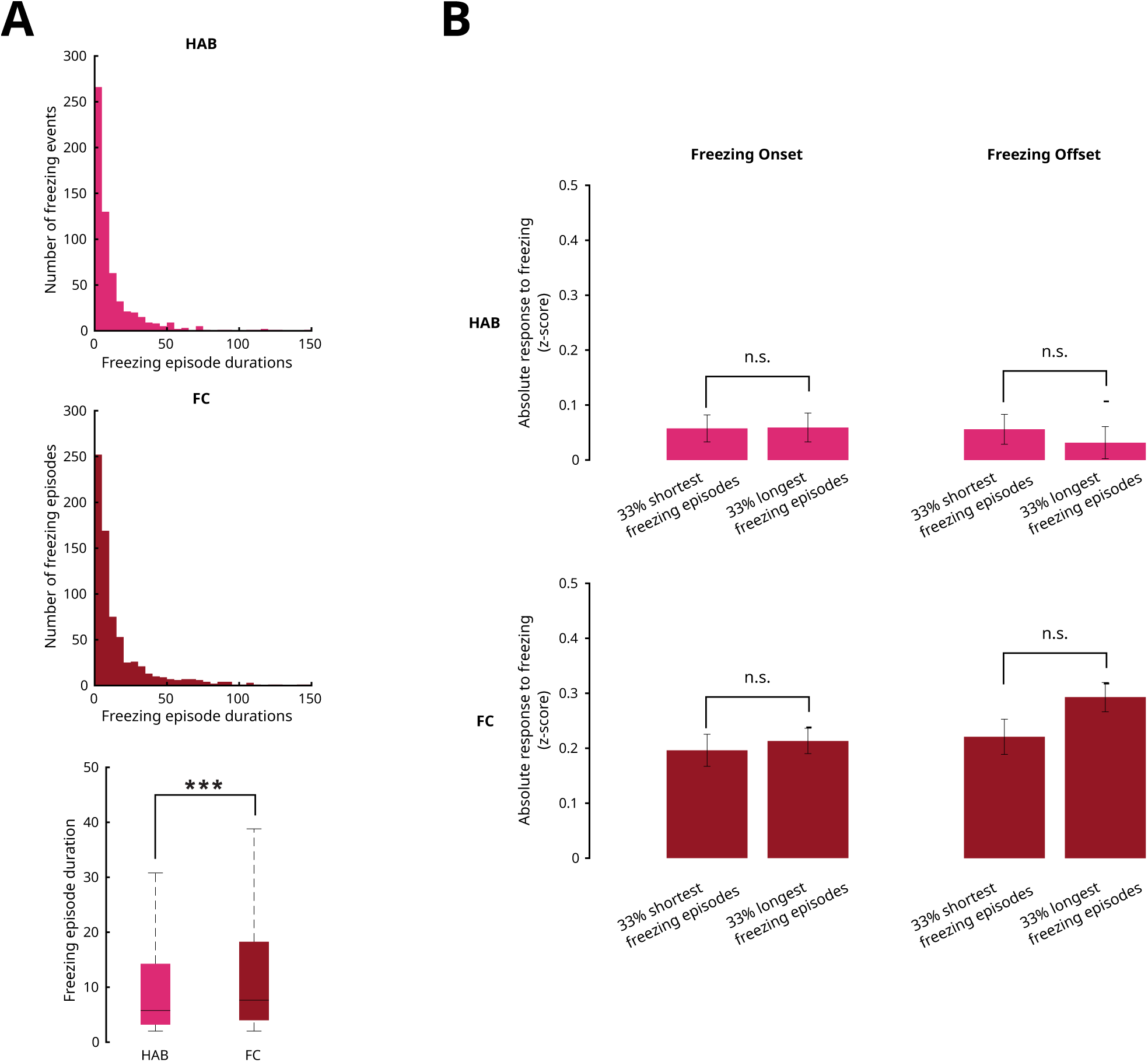
Neuronal responses to freezing is not affected by the duration of freezing episodes. (**A**) Distribution of freezing duration in HAB vs FC. (**B**) Comparison of the strength of the response (z-scored change in firing rate) to freezing for 33%of the shortest vs. 33% of the longest freezing episodes. The response to freezing is computed as in figure 2 (see Methods). In the case of negative responses, response magnitude was used for representation purposes (*p >* 0.05, Wilcoxon signed rank test).

**Supplementary Figure 7.**
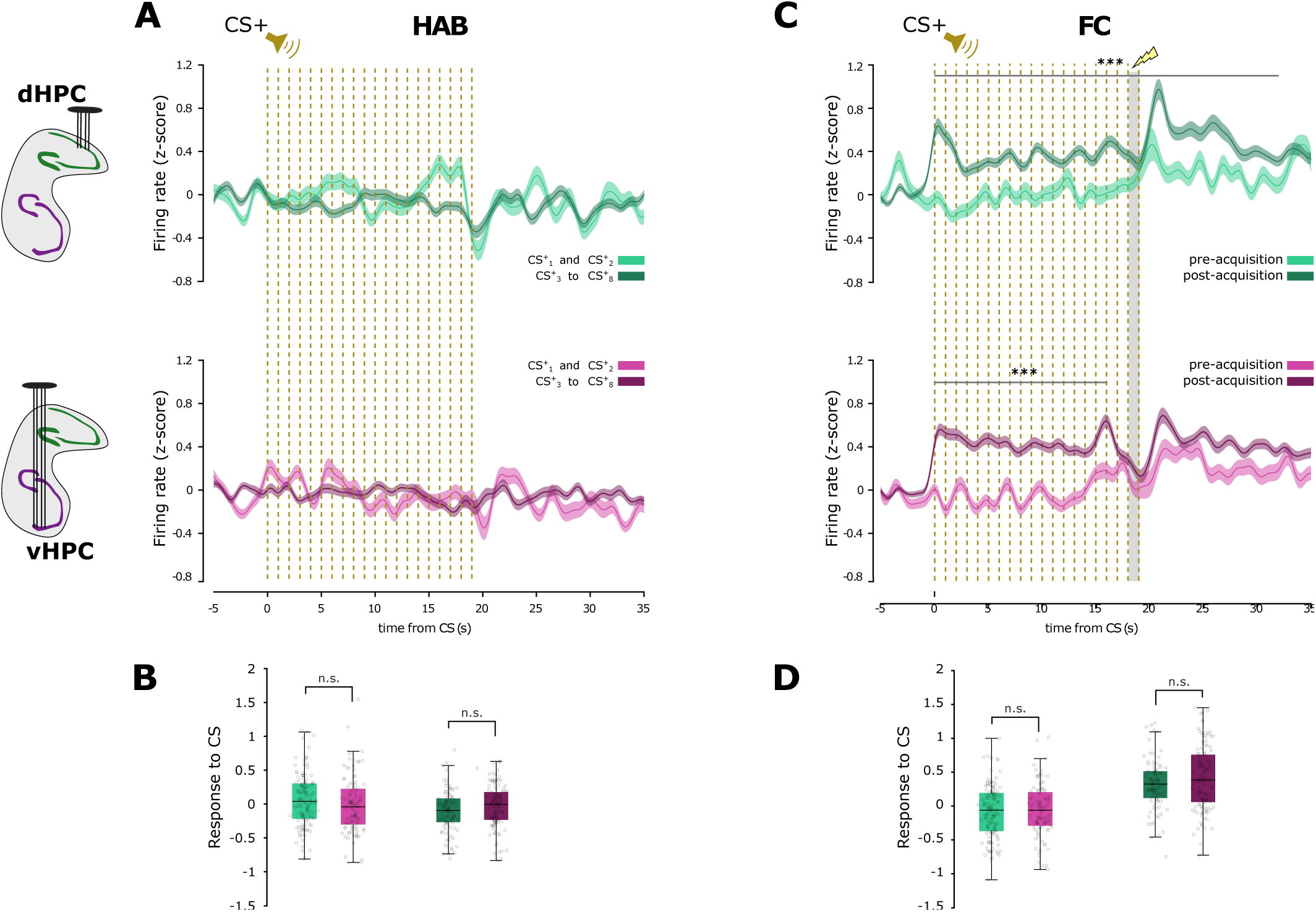
Non-speed-corrected firing rate responses to CS presentations. (**A**) Mean firing rate during the second habituation session for CS1 and CS2 (light curves) and CS3 to CS8 (dark curves) for dorsal hippocampal neurons (top) and ventral hippocampal neurons (bottom) (z-scored, baseline-subtracted but not speed-corrected). Same format as Figure 2G. (**B**) Comparison of the average responses to CS presentations (during the first 10 seconds after the CS) for CS 1-2 (top) and CS 3-8 (bottom) of dorsal (left) and ventral (right) hippocampal neurons. Both comparisons of dHPC vs vHPC are not significantly different. (*p >* 0.05, Wilcoxon signed rank test) (**C**,**D**). Same as A and B for FC sessions. Again, differences between dHPC and vHPC are not significant

**Supplementary Figure 8.**
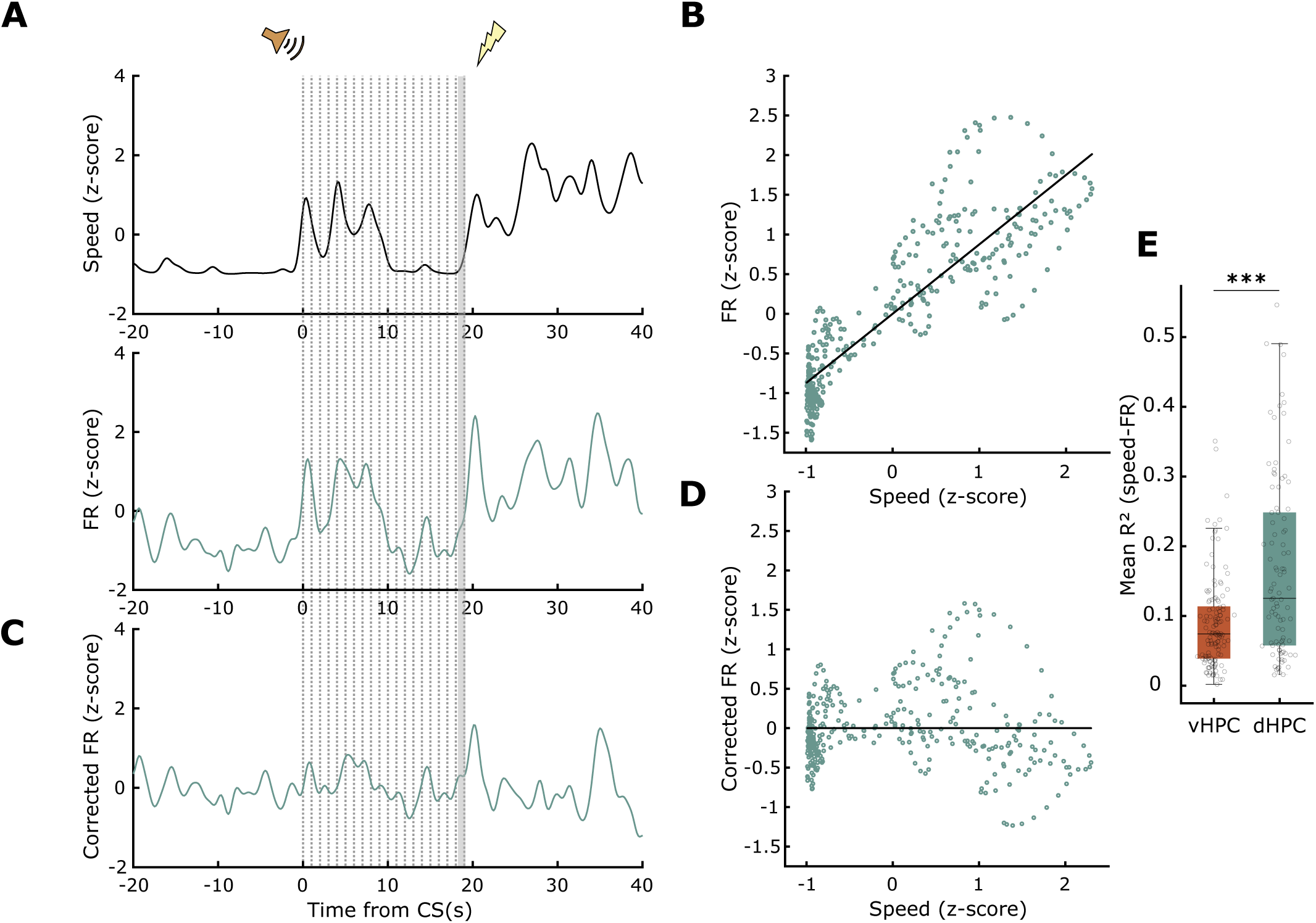
Correction for correlation between firing rate and speed. (**A**) Representative example of the speed (top) and firing rate of a dHPC neuron (bottom) during a CS presentation. (**B**) Linear regression for angular speed and firing rate for the same period as (A). The concentration of points at z-scored speeds equal to −1 correspond to moments where the animal was immobile. (**C**) Corrected firing rate for the example in (A) and (B). (**D**) Linear regression between speed and corrected firing rate. (**E**) Comparison of coefficients of determination for all dHPC and vHPC neurons computed from data from the same 1-minute periods (averaged over all 8 CSs during FC for each neuron).

**Supplementary Figure 9.**
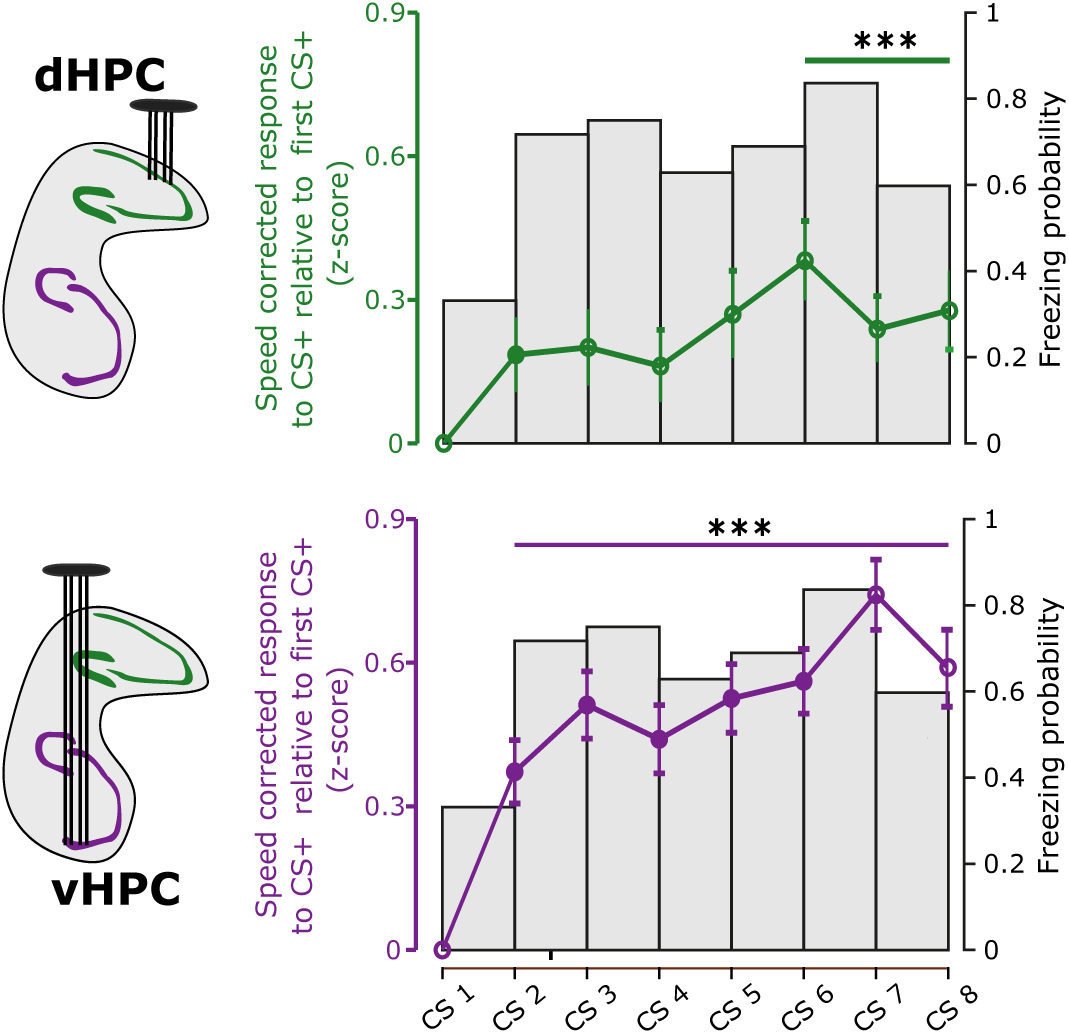
Coincidence of behavioral and neural responses. CS^+^ response changes relative to CS^+^1 (as baseline) compared with increase of freezing probability. Horizontal bars above curves show when the CS responses were significantly different from baseline (*p <* 0.001, Wilcoxon signed rank test).

**Supplementary Figure 10.**
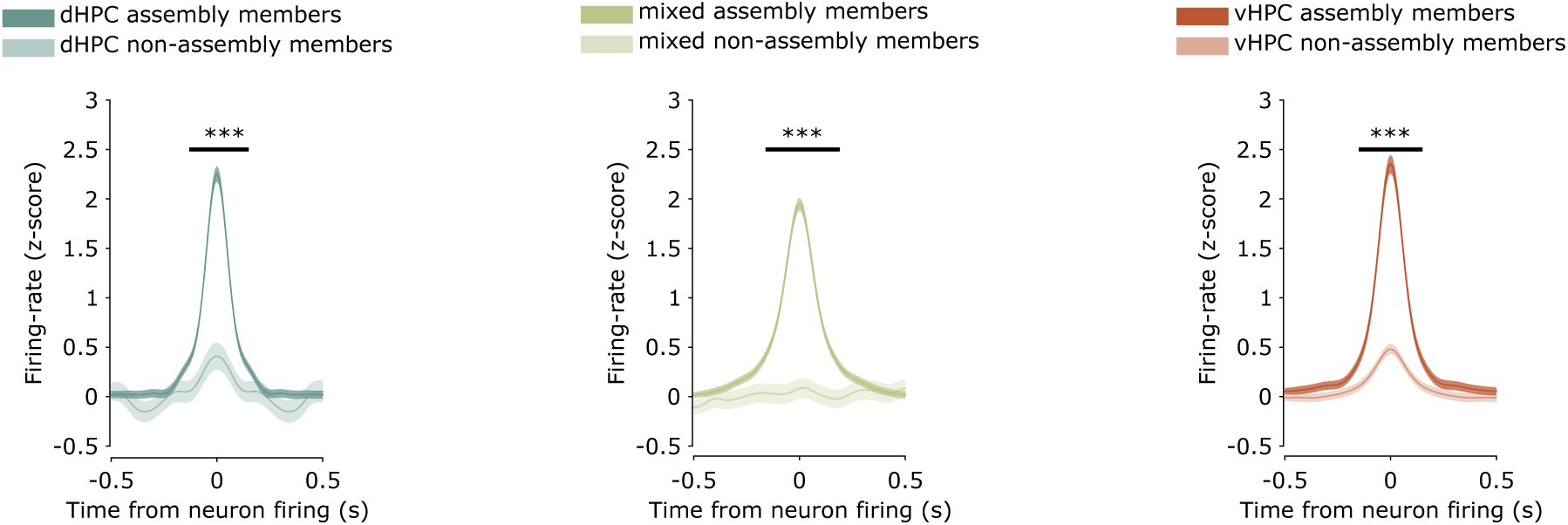
Cell assembly synchrony. Average cross-correlations of the activity of all neuron pairs that were both members of the same assembly vs. randomly selected pairs of neurons that were not members of the same assemblies. Horizontal bars above curves show when average cross-correlations of assembly members were significantly different from non-assembly members (*p <* 0.001, Wilcoxon ranksum test).

